# AlphaFold Protein Structure Database for Sequence-Independent Molecular Replacement

**DOI:** 10.1101/2021.09.10.459848

**Authors:** Lawrence Chai, Ping Zhu, Jin Chai, Changxu Pang, Babak Andi, Sean McSweeney, John Shanklin, Qun Liu

## Abstract

Crystallographic phasing recovers the phase information that is lost during a diffraction experiment. Molecular replacement is a dominant phasing method for the crystal structures in the protein data bank. In one form it uses a protein sequence to search a structure database for finding suitable templates for phasing. However, such sequence information is not always available such as when proteins are crystallized with unknown binding partner proteins or when the crystal is that of a contaminant. The recent development of AlphaFold has resulted in the availability of predicted protein structures for all proteins from twenty species. In this work, we tested whether AlphaFold-predicted *E. coli* protein structures were accurate enough for sequence-independent phasing of diffraction data from two crystallization contaminants for which we had not identified the protein. Using each of more than 4000 predicted structures as a search model, robust molecular replacement solutions were obtained which allowed the identification and structure determination of both structures, YncE and YadF. Our results advocate a general utility of AlphaFold-predicted structure database with respect to crystallographic phasing.

## 1. Introduction

Crystallographic phasing requires the retrieval of phase information that is lost during diffraction experiments. When there are no homology models, such phase information is recovered experimentally using isomorphous replacement preferably with their anomalous signals (Hendrickson, 2014, Liu & Hendrickson, 2015). With the accumulation of experimentally determined structures, molecular replacement (Rossmann, 1990, Evans & Mccoy, 2008) is becoming a routine method for crystallographic phasing. For example, 71% of deposited crystal structures in the PDB database (www.pdb.org), were determined using molecular replacement.

Molecular replacement exploits the similarity between known structures and the structure to be determined. Programs such as MOLREP (Vagin & Teplyakov, 1997), PHASER (Mccoy *et al*., 2007), and AMoRe (Navaza, 2001) have been developed. When protein sequence information is known, molecular replacement pipelines may be used to automate the process as implemented in MrBUMP (Keegan *et al*., 2018), BALBES (Long *et* al., 2008), and MRage (Bunkoczi *et al*., 2013). Using an *ab initio* modelling software such as ROSETTA (Das & Baker, 2008), predicted structures may also be used for molecular replacement as implemented in AMPLE (Bibby *et al*., 2012). However, there are cases in which the sequence information is unknown. Examples include the crystallization of contaminant proteins or unknown protein-binding partner proteins (Hungler *et al*., 2016). Under such scenarios, MoRDa (a non-redundant, annotated PDB database) (Vagin & Lebedev, 2015), ContaMiner/ContaBase (a collection of previously reported contaminant protein structures) (Hungler *et al*., 2016), and a SIMBAD pipeline (Sequence-Independent Molecular Replacement Based on Available Databases) (Simpkin *et al*., 2018) may be used for sequence-independent molecular replacement using database searching approaches. Among these tools, SIMBAD searches contaminant and MoRDa databases for a protein sequence-independent molecular-replacement (Simpkin *et al*., 2018).

Machine learning has been extensively used for protein structure predictions with the recent development of the revolutionary attention-based AlphaFold (Bouatta *et al*., 2021, Jumper *et al*., 2021) and RoseTTAFold algorithms (Baek *et al*., 2021). Both methods have enabled accurate prediction of protein structures approaching the fidelity of their crystal structures. In collaboration with an European Molecular Biology Laboratory (EMBL) team, AlphaFold released more than 350,000 predicted protein structures for twenty species including humans, and the predominant model systems including yeast, Arabidopsis, and *E. coli* (https://alphafold.ebi.ac.uk) (Tunyasuvunakool *et al*., 2021). The predicted structures cover all the coded proteins within each species. The AlphaFold-predicted structures may serve as a valuable new resource to support crystallographic phasing. It is therefore possible to use the species-wide structural databases for a protein sequence-independent molecular replacement for phasing diffraction data. This database approach may be of particular use for phasing proteins crystallized unexpectedly, proteolysis products, and structures of significant conformational changes. When crystallization of a protein with an unexpected binding partner protein, the AlphaFold database could be also used to identify potential binding proteins without the need for using mass spectrometry or protein sequencing.

During structural studies using X-ray crystallography, many proteins are expressed in *E. coli* and purified using various affinity columns. Often, in addition to protein of interest, *E. coli* contaminant proteins may bind either to the affinity resin or the protein of interest, and may co-purified and used for crystallization. Although crystallization of a contaminant protein is relatively rare, more contaminant structures have been identified as reported in the ContaBase database (Hungler *et al*., 2016). For new contaminant proteins it may take some effort to identify it through experimental phasing, mass spectrometry, protein sequencing, or using database searches. AlphaFold has generated a complete database of predicted structures for all folded protein sequences in *E. coli*. Here we sought to test whether this database supports crystallographic phasing in the absence of protein sequence information.

In our crystallization work on two plant proteins that were over-expressed in *E. coli*, we unexpectedly crystallized two contaminants and collected diffraction data to about 2.3-2.5 Å resolution. For one of them, we could not solve its structure using existing methods. In this work, we used the two contaminant data sets for sequence-independent molecular replacement. Using a relatively straightforward workflow, we show that AlphaFold-predicted structures can be used to phase both structures without any protein sequence information. Our work highlights the broad utility of the AlphaFold-predicted structure database for crystallographic analysis.

## 2. Methods

### 2.1 Sample preparation for YncE/P76116

*E. coli* contaminant protein YncE was co-purified while we worked on the expression of a plant Δ6 desaturase in BL21-Gold (DE3) cells (Novagen). The desaturase protein was over-expressed at 30 °C for 4 hours by addition of 0.2 mM IPTG to the cell culture with an A_600_ of 0.6. Harvested cells were re-suspended in resuspension buffer (30 mM MES, 33 mM HEPES, 33mM NaOAc pH 7.5) supplemented with 2 mM MgCl_2_ and 0.1 mg/ml DNase. The cells were lysed using a French Press and cell debris were removed by centrifugation at 25,000 x g for 30 min at 4°C. The clarified extract was loaded onto a Poros 20 HS column (Perceptive Biosystems, Framingham MA), washed with five column volumes of resuspension buffer, and eluted with a linear gradient of 0-1.2 M NaCl in the resuspension buffer. Desaturase fractions were pooled and concentrated, subjected to a size-exclusion HPLC column (TSKgel G3000SW column, Tosoh Bioscience), and eluted with 20 mM HEPES, pH 7.0, and 100 mM NaCl. The desaturase fractions were pooled and concentrated to 15 mg/ml.

Crystals were grown using the hanging drop vapor diffusion method consisting of 0.6 μl protein mixed with an equal volume of reservoir solution containing 0.2 M Li_2_SO4, 0.1 M MES, pH 6.0, and 20% PEG 4000. Plate-shaped crystals were flash-frozen with liquid nitrogen. Cryo-protectant was not added prior to freezing.

### 2.2 Sample preparation for YadF/P61517

*E. coli*. contaminant protein YadF was co-purified with the production of Arabidopsis Metacaspase 4 (AtMC4) in BL21 pLysS (DE3) cells (Novagen). Cells were lysed using a homogenizer and the soluble fraction of AtMC4 was collected for a three-step purification by nickel-nitrilotriacetic acid (Ni-NTA) affinity chromatography (HisTrap FF column, GE Healthcare, Inc.), ion exchange chromatography (HiTrap Q HP column, GE Healthcare, Inc.), and gel filtration (Superdex 200 10/300 GL column, GE Healthcare, Inc). Purified AtMC4 was then mixed and incubated with the excess molar amount of the inhibitor PPACK (Santa Cruz Biotechnology, Inc.). This mixture was further purified by gel filtration and the inhibitor-bound complex was concentrated to 8-10 mg/ml for crystallization.

Crystals were grown using the hanging drop vapor diffusion method. One μl of inhibitor-bound AtMC4 was mixed with an equal volume of precipitant that contains 100 mM sodium cacodylate, pH 6.8, and 1.8 M ammonium sulfate. For cryo-crystallography, crystals were transferred into the precipitant supplemented with 10% glycerol and were flash-cooled into liquid nitrogen for cryogenic data collection.

### 2.3 Diffraction data collection and reduction

Diffraction data were collected at the NSLS-II beamline FMX (17ID-2) at 100 K (Fuchs *et al*., 2016, Schneider *et al*., 2021). The beamline is equipped with an Eiger 16M detector. For YncE, we collected data at an X-ray wavelength of 0.979 Å. A total of 1800 frames were collected from a single YncE crystal with a rotation angle of 0.2°. For YadF, we collected data at an X-ray wavelength of 1.891 Å. A total of ∼1500 frames were collected from four YadF crystals with a rotation angle of 0.3°.

Single-crystal data sets were indexed and integrated independently using DIALS (Waterman et al., 2016) and then scaled and merged using CCP4 programs POINTLESS and AIMLESS (Evans *et al*., 2011, Evans & Murshudov, 2013) with the outlier rejection as implemented in PyMDA (Guo *et al*., 2018, Takemaru *et al*., 2020). For the YncE data, we rejected 700 radiation-damaged frames. For the YadF data, we rejected 948 radiation-damaged frames using a decay value of 1.0 as defined by frame_cutoff = [Min(SmRmerge) x (1+decay)], where Min(SmRmerge) is the lowest SmRmerge (reported in AIMLESS log file) within a single-crystal data set; and decay is a rejection ratio (Takemaru *et al*., 2020). The data collection and data processing statistics for the two data sets are shown in **Table 1**.

**Table 1.** Data collection and refinement statistics.

| **Data collection** | YadF/P61517 | YncE/P76116 |
| --- | --- | --- |
| Beamline | FMX (17-ID-2, NSLS-II) | FMX (17-ID-2, NSLS-II) |
| Wavelength (Å) | 1.891 | 0.979 |
| Space group | P4<sub>2</sub>2<sub>1</sub>2 | P2<sub>1</sub> |
| Cell dimensions a,b,c (Å) $\beta$ (°) | 67.52, 67.52, 85.25 | 53.17, 147.27, 96.90, 104.4 |
| Solvent content (%) | 43.0 | 51.8 |
| Bragg spacings (Å) | 36-2.3 (2.36-2.3) | 50-2.5 (2.56-2.5) |
| Total reflections | 222819 | 134117 |
| Unique reflections<sup>1</sup> | 9286 (665) | 47818 (3604) |
| Completeness (%) | 100.0 (100.0) | 95.9 (97.3) |
| $\langle I/\sigma(I) \rangle$ | 9.9 (2.2) | 7.3 (2.1) |
| R<sub>merge</sub> | 0.258 (0.912) | 0.087 (0.048) |
| Multiplicity | 24.0 (18.8) | 2.8 (2.8) |
| CC<sub>1/2</sub> (%) | 99.5 (81.2) | 98.1 (97.3) |
| **Refinement** |  |  |
| Resolution (Å) | 2.3 | 2.5 |
| No. reflections | 16710 | 87600 |
| R<sub>work</sub>/R<sub>free</sub> | 0.203/0.241 | 0.236/0.256 |
| No. atoms | 1756 | 10140 |
| Wilson B (Å<sup>2</sup>) | 31.0 | 30.7 |
| Average (Å<sup>2</sup>) | 40.2 | 46.6 |
| R.m.s deviations |  |  |
| Bond length (Å) | 0.002 | 0.002 |
| Bond angle (°) | 0.414 | 0.521 |
| PDB code | XXXX | XXXX |

<sup>1</sup>Values in parentheses are for the highest resolution range.

### 2.4 AlphaFold structures for database-driven molecular replacement

**Fig. 1** shows the workflow of using AlphaFold-predicted *E. coli* structure database for sequence-independent molecular replacement. From these twenty AlphaFold-predicted structure databases (https://alphafold.ebi.ac.uk/download), we downloaded all 4363 *E. coli* protein structures. Among these structures, we removed those with less than 50 residues from further use. Then, we set up a molecular replacement search using the remaining 4175 structures. For each structure, we performed molecular replacement in MOLREP (Vagin & Teplyakov, 1997) with both rotation and translation searches with a high-resolution data cut-off at 3.0 Å resolution. The structures displaying the highest rotation and translation peaks were used to narrow the molecular replacement search. For YncE, we removed 34 disordered residues from N-terminal and used MOLREP for multi-copy molecular replacement (Vagin & Teplyakov, 2000). For YadF, we tried molecular replacement in different space groups to find the one with the highest translation peak height.

**Figure 1.**
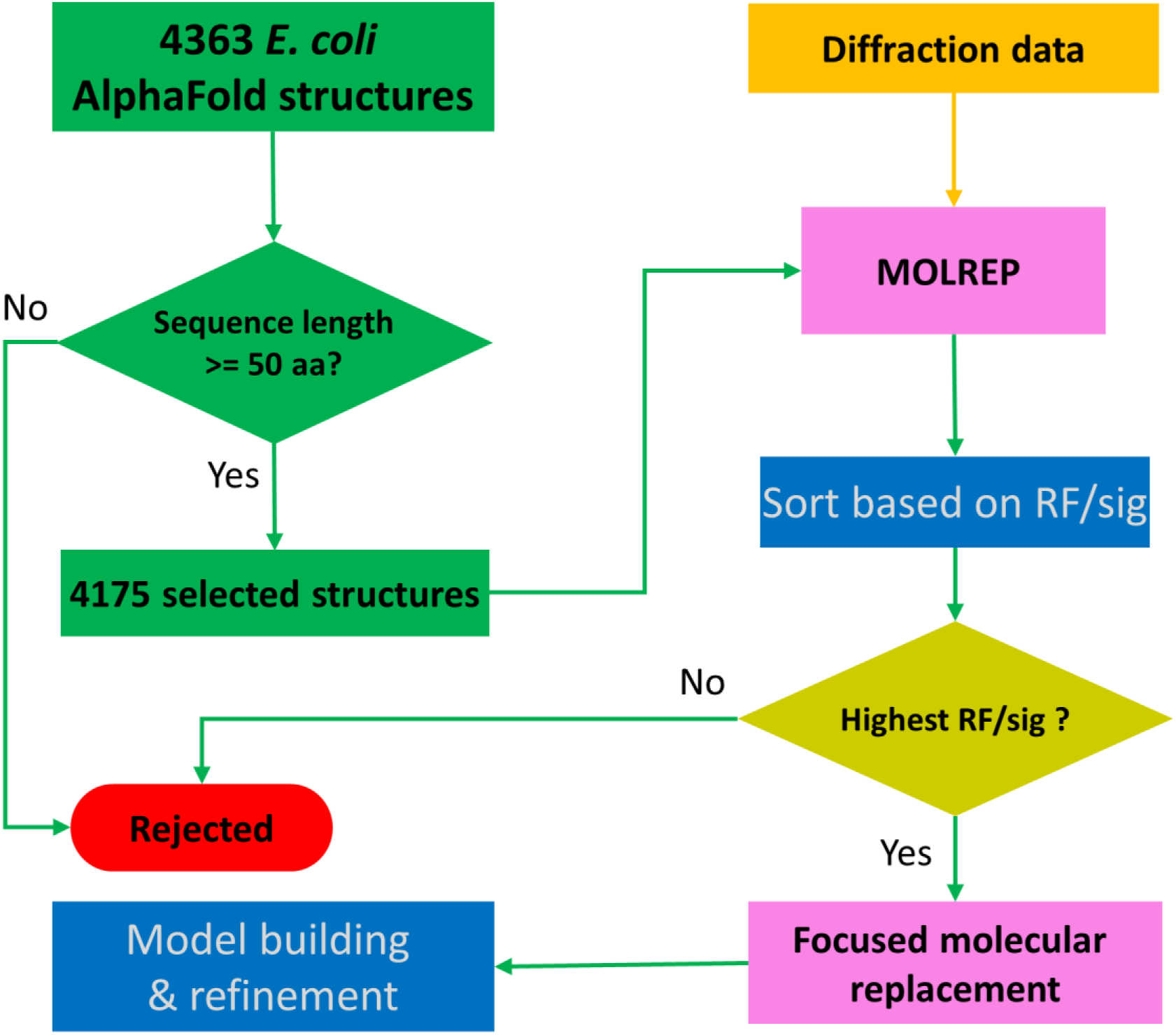
Schematic workflow of sequence-independent crystallographic phasing using AlphaFold-predicted *E. coli* structures. A total number of 4363 AlphaFold-predicted structures were downloaded from the AlphaFold structure database. After filtering based on protein sequence length, 4175 structures were selected for molecular replacement using MOLREP. The output candidate solutions were sorted based on TR/sig and the AlphaFold structure with the highest TR/sig peak height was selected for focused molecular replacement and downstream model building and refinement.

### 2.5 Model building and structure refinement

Iterative model building and refinement were performed in COOT (Emsley *et al*., 2010) and PHENIX.REFINE (Afonine *et al*., 2012, Echols *et al*., 2014), respectively. For the YncE data, Bijvoet pairs were averaged for structure refinement. For the YadF data, Bijvoet pairs were treated as two different reflections in structure refinement, and the resultant Fourier coefficients were used for calculation of Bijvoet-difference Fourier maps. We also used anomalous signals for a f” refinement (Liu *et al*., 2013) to find anomalous scattering elements in the YadF structure. For the f” refinement, the occupancies for the potassium and zinc ions were first estimated so that their refined individual B factors are close to the average B factors from their interacting protein and water molecules. We then refined f” in PHENIX.REFINE starting with f” values of zero for sulfur, potassium and zinc. The stereochemistry of the refined structures was validated with PROCHECK (Laskowski et al., 1993) and MolProbity (Chen et al., 2010) for quality assurance. The refinement statistics for the two data sets are shown in **Table 1**.

## 3. Results

### 3.1. AlphaFold structures for phasing YncE

During our work on the purification and crystallization of a plant desaturase, we co-purified YncE under crystallization conditions of 0.2 M Li_2_SO4, 0.1 M MES, pH 6.0, and 20% PEG 4000. We collected diffraction data and processed the data to *d*_min_ 2.5 Å in space group P2_1_ with unit dimensions a=53.2 Å, b=147.3 Å, c=96.9 Å, and β=104.4 °. Although the expected sequence identity for the desaturase to its homologous structures in PDB is beyond 80%, we were unable to solve the structure using the PDBs as search models, suggesting that this crystallized protein could be a contaminant. We used CCP4 online servers to search for contaminants but did not get a clear solution. To identify the contaminant, we also tried to repeat the crystallization and used mass spectrometry to identify the contaminant. Unfortunately, we were unable to reproduce the exact crystals.

With only the diffraction data available, we hypothesized that the contaminant protein must originate from *E. coli*. With the release of the AlphaFold-predicted *E. coli* structures, we reasoned that the crystallized contaminant should be represented in the AlphaFold structure database. We proceeded with the procedure described in **Fig. 1** to search for a monomer. All AlphaFold structures give their highest rotation and translation peaks beyond zero with a single structure, YncE (UNIPROT entry P76116), showed the highest RF/sig and TF/sig of 12.43 and 13.08, respectively (**Fig. 2a**).

**Figure 2.**
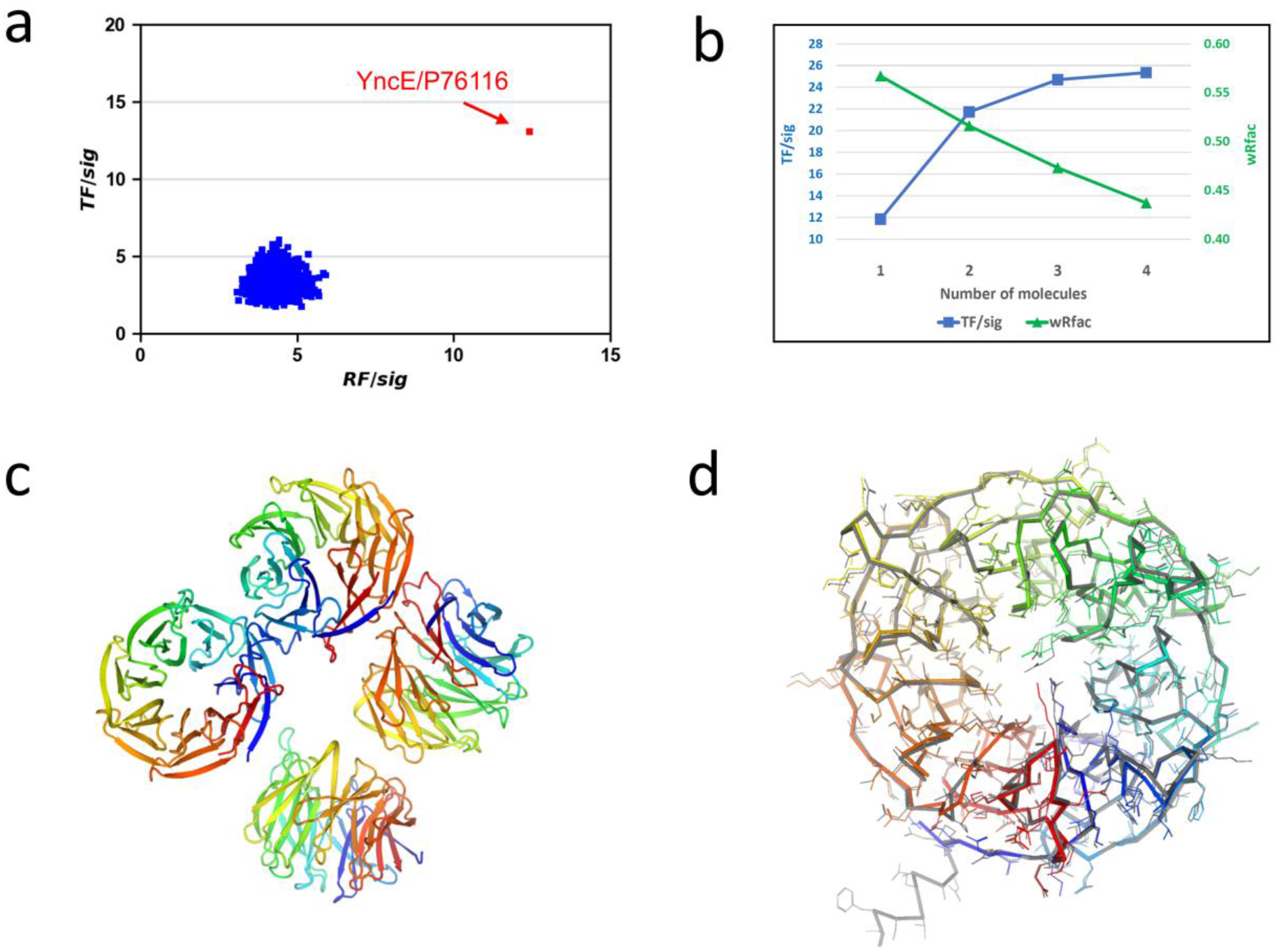
Alphafold structure for phasing *E. coli*. YncE. (**a**) Histogram of rotation and translation peaks. (**b**) Progressive molecular replacement while searching for four molecules in a.u. (**c**) Refined YncE structure. (**d**) Comparison of the refined structure with the AlphaFold structure.

Unit-cell content and self-rotation function analyses suggested the presence of multiple copies of YncE in the asymmetric unit (a.u.). We therefore performed focused molecular replacement searches for multiple copies using MOLREP. Visualization of the AlphaFold-predicted YncE structure indicated that it has a long N-terminal extension consisting of 34 poorly predicted/disordered residues. To assure that such a long extension would not affect the packing analysis in MOLREP, we removed the N-terminal 34 residues and used the truncated model for search of 2-5 monomers. We obtained the best results while searching for four monomers in a.u. and observed that both TF/sig and wRfac improved with an increasing number of monomers (**Fig. 2b**). With the four-monomer search, the final TF/sig and wRfac are 25.35 and 0.437, respectively, strongly indicating a correct solution for protein identification and structure determination.

The refined YncE structure has four molecules, each containing residues from 32 to 342 and forming a seven-bladed β-propeller structure (**Fig. 2c**). Except the N-terminal extension, the structure is very similar to the AlphaFold-predicted structure with an RMSD of 0.39 Å for 321 aligned C_α_ atoms (**Fig. 2d**). However, we found that many side chains have different conformations, perhaps due to crystal contacts or disordered conformations.

In the UNIPROT entry for P76116/YncE, two PDBs (3VGZ and 3VH0) are reported, one crystallized in C222_1_ lattice and the other crystallized in I4_1_ lattice (Kagawa *et al*., 2011). Our P2_1_-form structure is a new contaminant structure. The P2_1_-form structure has an RMSD of 0.44 Å with the C222_1_-form structure and 0.37 Å with the I4_1_-form structure, indicating that all three structures are very similar although being crystallized in different space groups. **Table 2** summarizes the detailed crystallographic comparison of the YncE structure determined in three different lattices.

**Table 2.**
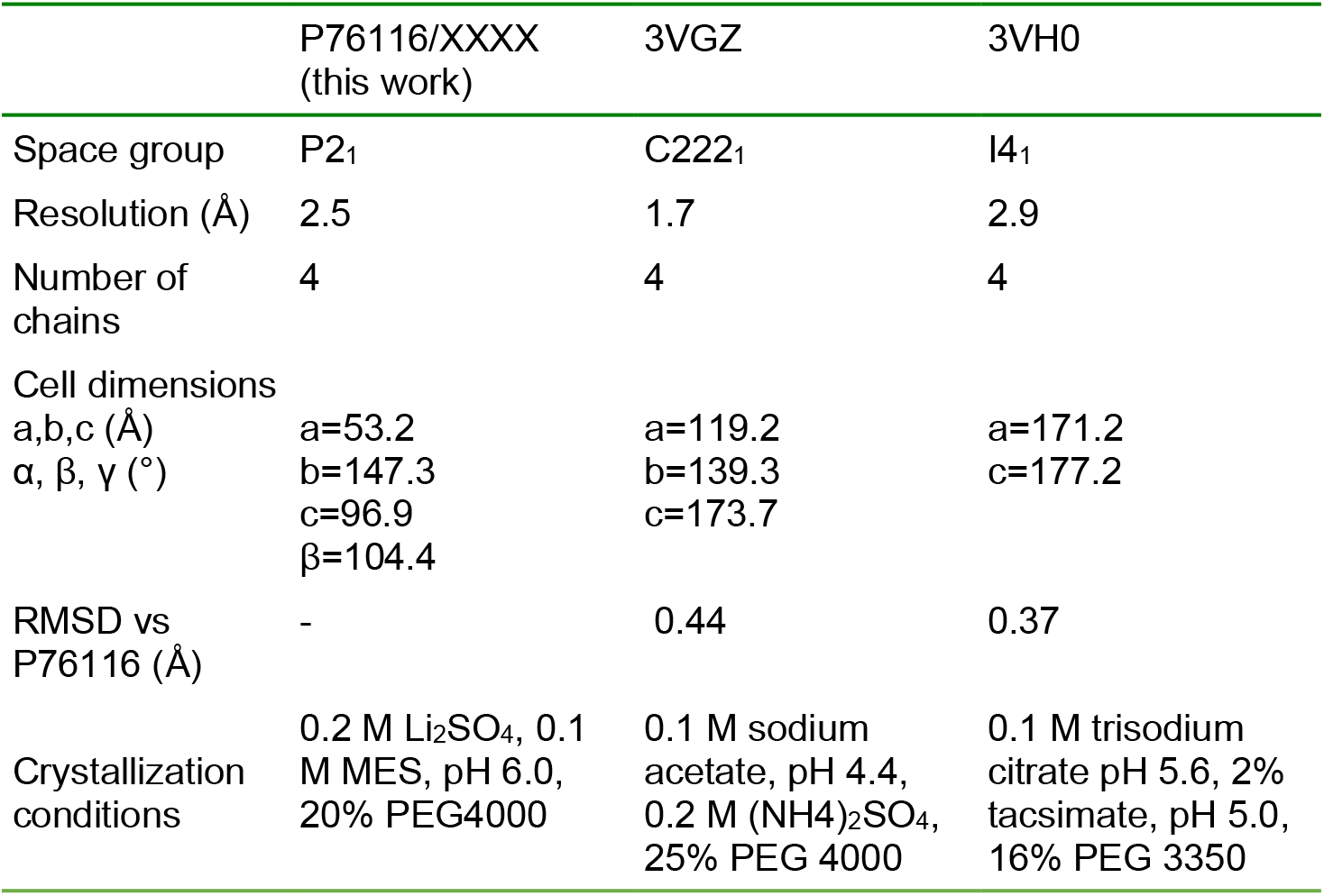
Comparison of YncE/P76116 structure with PDB structures listed under UNIPROT entry P76116.

### 3.2 AlphaFold structures for phasing YadF

*E. coli* YadF is another contaminant protein that was co-purified with an Arabidopsis metacaspase 4 (AtMC4). AtMC4 is a cysteine protease and we have previously determined its structure in an apo form (Zhu *et al*., 2020). To get a complex structure of AtMC4 with a protease inhibitor PPACK, we attempted to crystallize the complex for structural analysis. Crystals with dimensions of about 20-30 μm were obtained under the crystallization conditions of 0.1 M sodium cacodylate, pH 6.8 and 1.8 M (NH4)_2_SO_4_. We collected diffraction data from four crystals at a relatively longer X-ray wavelength of 1.891 Å. The processed data at d_min_ 2.3 Å has a tetragonal lattice with unit-cell dimensions of a=67.5 Å and c=85.3 Å. However, we couldn’t solve its structure using the AtMC4 structures of either the full-length or its truncations. Therefore, we suspected that this could be another *E. coli* contaminant and may be suitable for structure determination through searching the AlphaFold-predicted structure database.

Using the same workflow described above for YncE, we performed molecular replacement searches using MOLREP for each of the 4175 structures. **Fig. 3a** shows the histogram plot for RF/sig and TF/sig. Although there are four targets with highest translation peaks beyond 10 (UNIPROT entries P0CF69, P75971, P0CF68, and P61517), P61517/YadF is the only target with the highest rotation peak at 9.04, suggesting it is a possible solution for downstream model building and refinement. YadF has 220 residues, the unit-cell content analysis suggested a single molecule in a.u. with an estimated solvent content of 43%. The initial refinement in PHENIX.REFINE yield an R/free R of 0.30/0.39, suggesting larger structural differences relative to the AlphaFold-predicted structure. Therefore, we rebuilt the model using ARP/WARP (Langer *et al*., 2008). ARP/WARP produced a nearly complete model of 208 residues in one chain with an R/free R of 0.194/0.252, indicating a correct identification and structure determination using the AlphaFold structure database.

**Figure 3.**
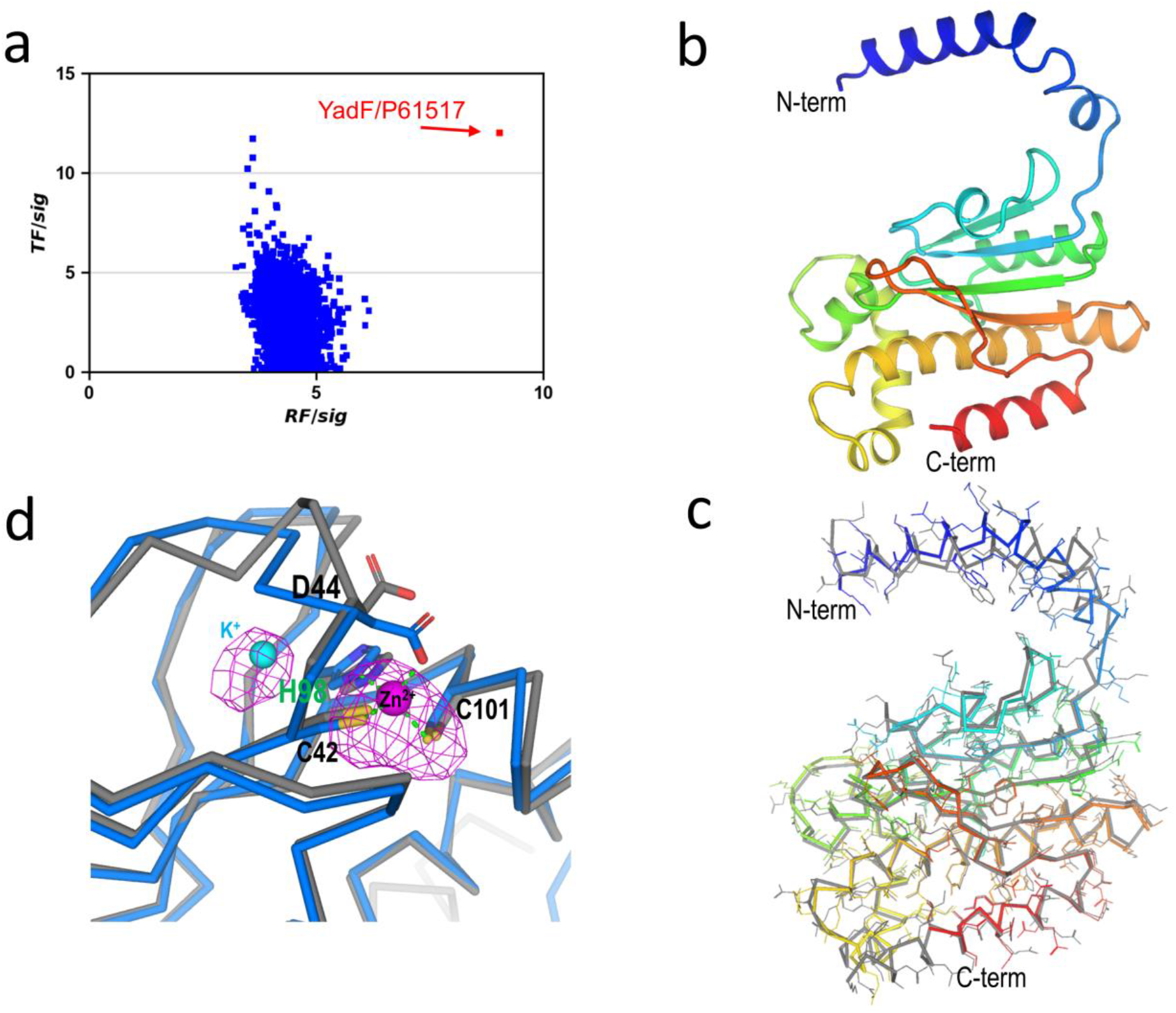
AlphaFold structure for phasing *E. coli*. YadF. (**a**) Histogram of rotation and translation peaks. (**b**) Refined YadF structure. (**c**) Comparisons with the AlphaFold structure. (**d**) Active-site structure. Residues interacting with the zinc site are shown as sticks. Bijvoet difference Fourier map for anomalous scatterers were shown as magenta isomeshes contoured at 3σ. As a comparison, AlphaFold-predicted structure is shown in gray.

The refined structure has 211 residues, and its structure is shown in **Fig. 3b**. The structure has an N-terminal α-helix domain and a C-terminal mixed αβ domain. Compared with the AlphaFold-predicted structure, the RMSD is 1.18 Å for 206 aligned C_α_ atoms. Most structural differences are on the N-terminal helix and the loop connecting it to the αβ domain (**Fig. 3c**).

YadF is a carbonic anhydrase whose activity is zinc dependent (Cronk *et al*., 2001). We had collected data at an X-ray wavelength of 1.891 Å at which the theoretical anomalous signal f″ is 0.98 e. Therefore, we used an f″ refinement to characterize zinc anomalous signals (Liu *et al*., 2013). With an estimated occupancy of 1.0 for the zinc site, the refined f″ is 0.94 e, clearly validating the specialization of the zinc site. Zinc is coordinated with two cysteine residues (Cys42 and Cys101), His98, and Asp44. **Fig. 3d** shows the Bijvoet difference Fourier densities for the active site. The Bijvoet densities cover zinc as well as two sulfur atoms. Surprisingly, next to the zinc/sulfur densities, we observed an extra electron density next to His98. To identify the type of anomalous scatterers associated with this density, we performed the f″ refinement with a candidate ion of Zn^2+^, Ca^2+^, K^+^, or Na^+^. Through the f” refinements, the only reasonable fit for this anomalous scatterer is K^+^ with an occupancy of 0.6 and a B-factor of 33.5 Å^2^. However, we did not include K^+^ either in protein purification or crystallization. It’s exact origin and potential functional role will therefore be the subject of further investigation.

The AlphaFold-predicted structure does not contain any ions, neither Zn^2+^ nor K^+^. Structural superimposition of the AlphaFold structure with the ion-bound YadF structure indicates conformational changes of Asp44 (**Fig. 3d**). Interestingly, the same residue has been proposed to undergo conformational change so that substrate CO_2_ can approach Zn^2+^ to form a CO-Zn^2+^ species (Cronk *et al*., 2001). Thus, it is possible that the AlphaFold-predicted structure might resemble an intermediate state of YadF, at least for the active site structure.

Under UNIPROT entry P61517, there are four reported PDBs (1I6O, 1I6P, 1T75, and 2ESF) (Cronk *et al*., 2001, Cronk *et al*., 2006), all determined in tetragonal lattices but with different crystallization conditions. Our structure has a RMSD between 0.35 and 0.79 Å compared to these structures. **Table 3** summarizes detailed crystallographic comparison of YadF under different crystallization conditions.

**Table 3.**
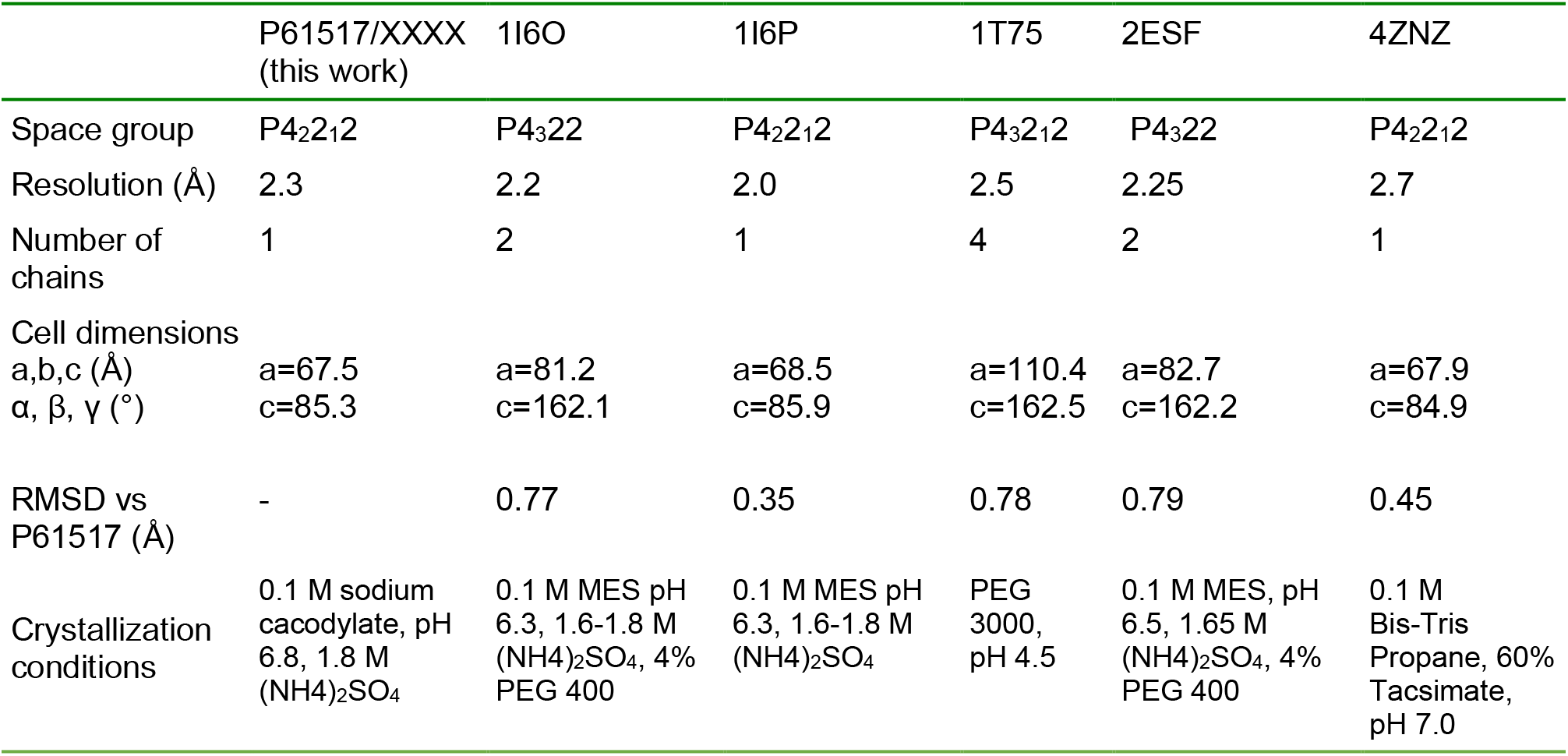
Comparison of YadF/P61517 structures with PDB structures listed under UNIPROT entry P61517.

## 4. Discussion

### 4.1 AlphaFold-predicted structure database

Crystallizing protein contaminants is a relatively common problem. In this work we demonstrate that AlphaFold-predicted *E. coli* structures can be useful for molecular replacement to identify unknown crystallized contaminant proteins and determine their structures. In our tests, we did not modify the predicted structures for the initial molecular replacement searches even though these predicted structures may contain unstructured extensions and poorly predicted regions such as we found with the N-terminal long extension in YncE.

For the two contaminant structures that we determined using AlphaFold-predicted structure database, YncE is a new contaminant. Although there are two crystal structures (PDB entries 3VGZ and 3VH0), we did not get a clear solution while trying database search approaches using the CCP4 online server. As a comparison, for YadF, in addition to using AlphaFold structure database, we can find a solution using its unit cell dimensions to search the PDB database, and PDBs 1I6P and 4ZNZ were identified. It turned out that PDB 4ZNZ was already reported as a crystallization contaminant (Niedzialkowska *et al*., 2016) but crystallized in a different condition (**Table 3**). It is noted that the YadF structure in this work has a larger RMSD with the AlphaFold-predicted structure (1.2 Å) than with other crystal structures (0.35 – 0.79 Å). As shown in **Fig 3c**, largest structural differences are located at the N-terminal helix. In YadF crystal structures, this helix is stabilized by forming a dimer with its symmetry mate (Cronk *et al*., 2001). As a contrast, AlphaFold-predicted structure is a monomer, and the N-terminal helix can thus be more flexible.

Phasing with an *E. coli*. structure database has multiple advantageous over the PDB database. First, the predicted structures contain only single-chain structures, which may be used directly for rotation search with no need for further processing, such as removing non-protein components or splitting a protein complex into individual components. Secondly, the predicted structure is based on the entire encoded protein sequences. Consequently, using such a database gives a higher chance to find a promising and suitable structure template for phasing. Although in this work we used *E. coli* structures for identification and determination of contaminant structures, AlphaFold has predicted 350,000 structures for proteins from 20 species (Tunyasuvunakool *et al*., 2021); and those structure databases may be well suited for phasing contaminant structures from other expression hosts such as mammalian cells, yeast, Arabidopsis and etc. Thirdly, AlphaFold structures may be used to identify and phase unexpected proteolytic fragments or unexpected binding partner proteins.

Using a domain-structure database and modelled structure for phasing has been previously implemented in MoRDa and AMPLE, respectively (Bibby *et al*., 2014, Vagin & Lebedev, 2015). However, due to the limited number of structural domains and the inaccuracy associated with the modelling, database-based phasing is not routine, and is normally used as a method of last resort after exhausting other phasing strategy options. As AlphaFold-predicted structures have an improved accuracy relative to experimental structures, molecular replacement using AlphaFold structures could have more routine applications even for *de novo* phasing for which there is no homologous structure. The AlphaFold algorithm uses an artificial intelligence model that was extensively trained with available PDBs and sequence databases (Jumper *et al*., 2021). Hence the AlphaFold-predicted structures could be biased toward known structures. Accordingly, novel protein structures with novel folds are needed to improve the prediction accuracy of AlphaFold. Based on our findings, we speculate that an increasing number of crystal structures will be phased using AlphaFold-predicted structure workflows.

### 4.2 Combining AlphaFold phasing with anomalous signals

Perhaps due to the existence of prior crystal structures for both YncE and YadF, AlphaFold-predicted structures are quite accurate, with a RMSD of 0.30 Å and 1.18 Å, to their refined structures (**Fig. 2d, 3c**). When there are only remote or no homologous structures, AlphaFold-predicted structures may be not sufficient for phasing solely through molecular replacement. We propose that molecular replacement with anomalous signals, e.g. MR-SAD (Panjikar *et al*., 2009), might be a highly productive option.

For YadF, we collected long-wavelength data at 1.891 Å which allowed the characterization of anomalous scatters zinc, potassium, and sulfur atoms within the structure. To see whether anomalous signals will enhance AlphaFold-based crystallographic phasing, we tested MR-SAD (Panjikar *et al*., 2009) using the PHASER_EP pipeline (Mccoy *et al*., 2007). With the initial phases from the AlphaFold structure, PHASER_EP identified seven anomalous scatterers with a figure-of-merit of 0.467. The MR-SAD map is of high quality; the pipeline can build 201 residues in eight fragments, with the longest fragment representing 71 residues. Subsequently, ARP/wARP can build the exact model as starting from the AlphaFold structure without using anomalous signals. For YadF, anomalous signals did not help much because ARP/wARP overcame the model errors (for example the Nterminal helix, **Fig. 3c**) through automated model building. For cases where the model is not accurate enough or the diffraction data are not of high enough resolution, MR-SAD may help to solve structures that are otherwise very challenging or even currently considered unsolvable. Most proteins contain intrinsic sulfur atoms that are native anomalous scatters of long-wavelength X-rays. So, to optimize the use of AlphaFold-predicted structures for phasing a *de novo* structure, it might be advantageous to collect long-wavelength native-SAD data preferably using a helium flight path if available. Then these anomalous signals from sulfur atoms can be used for AlphaFold-based phasing using MR-SAD.

## 5. Concluding remarks

Using the AlphaFold-predicted *E. coli* structure database, we identified and determined structures for two crystallization contaminants without protein sequence information. The molecular replacement solutions and the structural comparison of refined structures with those AlphaFold-predicted structures suggest that the predicted structures are of high accuracy for crystallographic phasing and will likely be integrated into other structure determination pipelines.

## Acknowledgements

This work was supported in part by Brookhaven National Laboratory LDRD 22-008 and NIH grant GM107462. Q. L. was supported by the U.S. Department of Energy, Office of Science, Office of Biological and Environmental Research, as part of the Quantitative Plant Science Initiative at BNL. J.C. and J.S. were supported by Division of Chemical Sciences, Geosciences, and Biosciences, Office of Basic Energy Sciences, United States Department of Energy Grant DOE KC0304000. The FMX beamline is part of the Center for BioMolecular Structure (CBMS) which is primarily supported by the National Institutes of Health, National Institute of General Medical Sciences (NIGMS) through a Center Core P30 Grant (P30GM133893), and by the DOE Office of Biological and Environmental Research (KP1607011). NSLS-II is supported in part by the U.S. Department of Energy, Office of Science, Office of Basic Energy Sciences Program under contract number DE-SC0012704 (KC0401040).

## Data availability

Atomic coordinates and structure factor files have been deposited in the RCSB Protein Data Bank (PDB) under the accession codes XXXX and XXXX.

